# What ecological factors favor parthenogenesis over sexual reproduction? A study on the facultatively parthenogenetic mayfly *Alainites muticus* in natural populations

**DOI:** 10.1101/2021.09.10.459770

**Authors:** Maud Liegeois, Michel Sartori, Tanja Schwander

**Affiliations:** Department of Ecology and Evolution, University of Lausanne, CH-1015 Lausanne, Switzerland; Cantonal Museum of Zoology, Palais de Rumine, Place de la Riponne 6, CH-1014 Lausanne, Switzerland

**Keywords:** sex ratios, facultative parthenogenesis, mate limitation, natural populations, ecological factors

## Abstract

Different reproductive modes are characterized by costs and benefits which often depend on ecological contexts. Benefits of sex are expected to increase under complex biotic interactions, whereas parthenogenesis can be beneficial for reproductive assurance when females are mate limited. Here, we study how different ecological contexts influence the frequency of sex or parthenogenesis in the facultatively parthenogenetic mayfly *Alainites muticus*. We first verified that high parthenogenetic frequencies translate into female-biased population sex ratios. We then measured the population sex ratio (a proxy for parthenogenetic capacities), density of individuals (mate limitation) and community diversity (biotic interaction complexity) for 159 *A. muticus* populations, and used structural equation modeling to investigate their direct and indirect influences on sex ratios. We found no effect of community diversity or altitude on sex ratios. Furthermore, even when females can reproduce parthenogenetically, they generally reproduce sexually, indicating that the benefits of sex exceed its costs in most situations. Sex ratios become female-biased in low population densities, as expected if mate limitation selects for parthenogenesis. Mate limitation might be widespread in mayflies because of their very short adult lifespan and limited dispersal, which can generate strong selection for reproductive assurance and may provide a steppingstone towards obligate parthenogenesis.

## Introduction

Sexual reproduction is by far the most abundant reproductive mode in the animal kingdom, a pattern that is difficult to explain given the many costs associated with sex (reviewed in Lehtonen *et al*., 2012). A number of theories have been developed that can help explain the advantage of sex, but experimental tests in natural populations remain scarce (Neiman *et al*., 2018). An ideal approach to identifying conditions under which sex provides benefits is to study variations in the frequency of sex in species capable of facultative parthenogenesis. Such species avoid problems inherent to comparisons between sexual and parthenogenetic species, as there may be species-specific traits that are confounded with reproductive mode. Facultative parthenogenesis is rare among animals overall, but is widespread among mayflies (Ephemeroptera), with at least 49 species reproducing by facultative parthenogenesis (Liegeois *et al*., 2021). Here, we used the facultatively parthenogenetic mayfly species *Alainites muticus* (Baetidae) to identify ecological correlates of the frequency of sex *versus* parthenogenesis in natural populations.

Specifically, we evaluated three ecological contexts hypothesized to affect the costs and benefits of sex *versus* parthenogenesis. First, population densities can affect the level of mate availability, and parthenogenesis can provide benefits through reproductive assurance, especially when males are scarce (Gerritsen, 1980; Schwander *et al*., 2010). Population densities, by affecting the frequency of harassment by males, can also favor parthenogenesis at low densities because in this case the cost of resisting male harassment is low (Gerber & Kokko, 2016). Second, because biotic interactions are the drivers of co-evolutionary dynamics (reviewed in Lively & Morran, 2014), sexual reproduction can be favored in complex communities with various types of species interactions and high levels of competition (“the Red Queen hypothesis”, *e.g*., Van Valen, 1973; Jaenike, 1977; Hamilton, 1980). Similarly, the Tangled Bank and Sib Competition hypotheses propose explanations for the prevalence of sex in relatively saturated communities, where competition for resources is likely to be continuous and intense (*e.g*., Williams, 1975; Bell, 1982; Scheu & Drossel, 2007).

Increasing benefits of sex in complex communities is, for example, believed to explain why parthenogenetic species are rare in the tropics (Bell, 1982). Third, a frequently observed pattern in geographical parthenogenesis is that obligately asexual species are more likely to be found at higher altitudes than their sexual counterparts, for different but often unknown reasons (reviewed in Tilquin & Kokko, 2016). However, there are currently no data available to test whether a similar pattern of geographical parthenogenesis holds for facultative parthenogens.

In order to elucidate how biotic and abiotic ecological variables, such as the density of individuals, community complexity, and altitude, can affect reproductive strategies, we used structural equation modeling (SEM), a multivariate statistical approach that allows considering interactions between factors and hidden indirect effects (*e.g*., Grace *et al*., 2010; Eisenhauer *et al*., 2015; Fan *et al*., 2016). We thus tested, using 159 populations of *Alainites muticus*, whether low population densities favor parthenogenesis, complex communities favor sex, the frequency of parthenogenesis increases with altitude, and how these variables are interconnected with each other.

We focused on the mayfly species *A. muticus* (Baetidae) because of its widespread distribution across altitudinal ranges, and its ability to reproduce by facultative, female-producing parthenogenesis (thelytoky; Degrange, 1960). In order to estimate the frequency of sex and parthenogenesis in each population, we used population sex ratios as a proxy. With facultative parthenogenesis, population sex ratios are expected to be female-biased if females reproduce at least partly by parthenogenesis, whereas population sex ratios are expected to be approximately equal if females reproduce sexually. To corroborate that population sex ratios indeed reflect the capacity for parthenogenesis, we measured the hatching success of unfertilized eggs laid by females and tested whether hatching successes were correlated with population sex ratios.

Finally, we also assessed the stability of population sex ratios over time, as well as the temporal variation of population densities and community diversities, by integrating information from two surveys per population, separated by five years.

## Material & Methods

### Study sites, sex ratios and ecological data

We used samples of *Alainites muticus* from 159 sites (*i.e*., populations). These samples stem from a biodiversity monitoring survey performed at 500 sites across Switzerland, following the IBCH method (described in Stucki, 2010). This standardized sampling technique is designed to provide an indication of the water quality of streams and rivers on the basis of aquatic macro-invertebrate communities. In brief, each site is sampled from early-March to mid-June, covering the ecologically relevant altitudinal gradient in Switzerland (from 193 to 2635m, sampling at higher altitudes is done later in the season). In order to cover all substrates of the riverbed and the range of current velocity, eight square areas (25 × 25 cm) are sampled by scraping the riverbed during 30 seconds and by capturing the macro-invertebrates using a standard-sized net. The corresponding samples are then stored in 80% ethanol. Individuals are sorted to the family level and used for the calculation of a water quality index. For each sampling site, standard data are systematically collected (*e.g*., altitudes, GPS coordinates, etc.). Nymphs of three insect orders (Ephemeroptera, Plecoptera and Trichoptera) are further identified to the species level and stored at the Museum of Zoology in Lausanne, from where we obtained the *A. muticus* samples. The biodiversity monitoring started in 2010 and it took five years to cover all 500 sites across Switzerland. Since 2015, each site has been sampled a second time in order to evaluate how water quality changes over time.

*Alainites muticus* occurred at 214 of the 500 sites across an altitudinal range of 195-1931m, and for further analysis, we selected all sites where more than 15 *A. muticus* individuals were present (n = 159/214, Fig. 1). For each site, we first quantified sex ratios by sorting male and female nymphs. Males are easily identifiable in early instars by their large eyes and their external genitalia. Nymphs smaller than two millimeters were excluded from sex ratio calculations, as they cannot be sexed reliably. We then assessed biotic and abiotic variables of each site: population densities, community diversities and altitudes. We used two different density estimates, the total number of nymphs found in each population, and the sexed individuals only, but results were not affected by the estimate used. Community diversities were estimated using the Shannon’s index (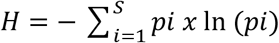, where *p_i_* is the proportion of individuals belonging to the *i*th species in the dataset of interest). Given the different level of identification for different taxa, (species level for Ephemeroptera, Plecoptera and Trichoptera, family level for all other macro-invertebrates), the index was based on abundances at the family level. Note that we also used other biodiversity indexes including Margalef’s index, Simpson’s index and Hill’s numbers (Simpson 1949; Margalef 1956; Heip *et al*., 1988; Magurran 2004; Keylock 2005; Jost 2006; Daly *et al*., 2018) in order to evaluate whether our conclusions were consistent according to different diversity measures.

In order to test whether *A. muticus* sex ratios, densities and macro-invertebrate community diversities were relatively stable or fluctuate over time, we used the 92 of the 159 sites which were surveyed twice (with a time gap of five years). For the remaining 67 populations, we only have data from a single sampling event.

**Figure 1.**
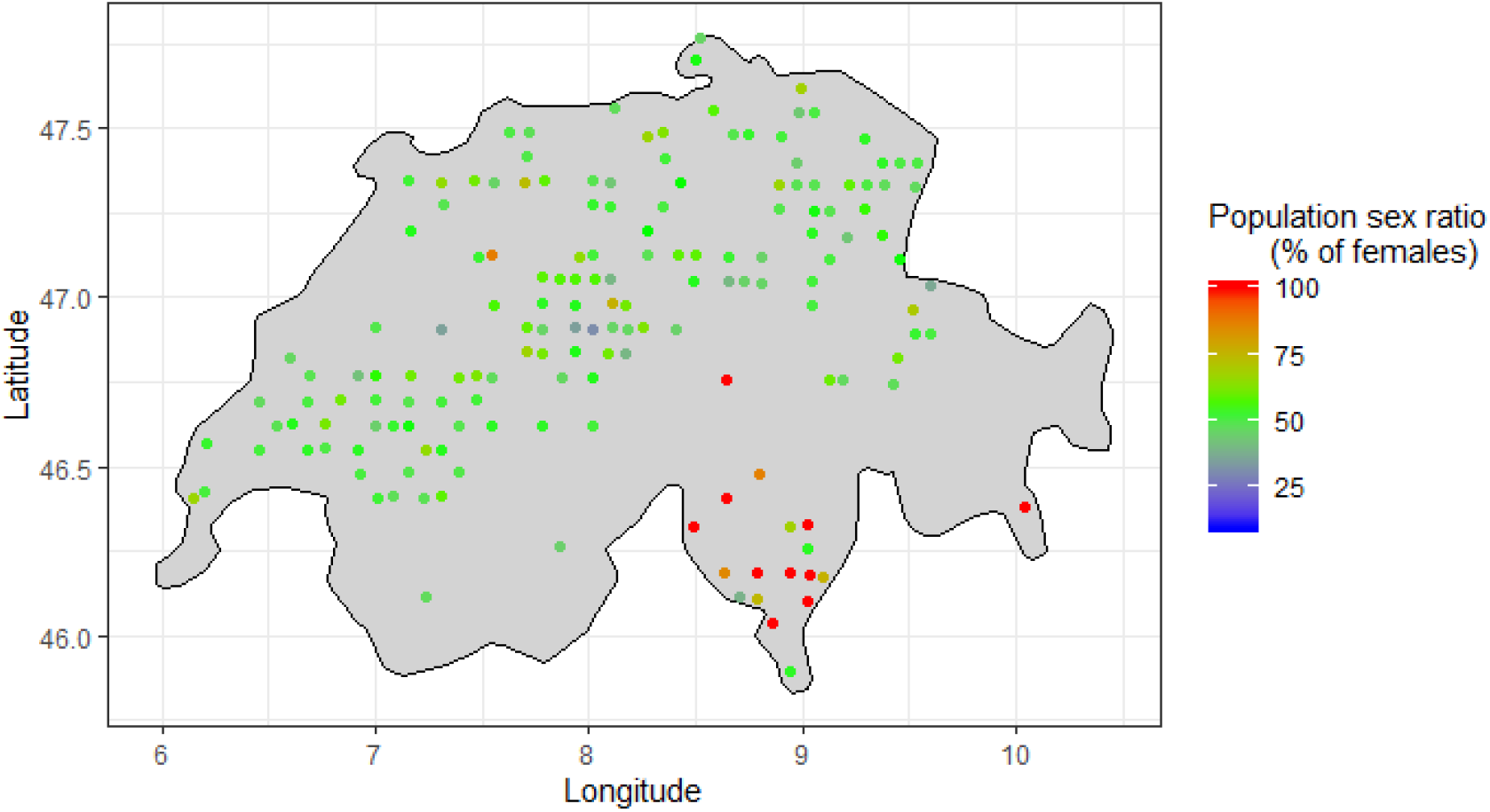
Distribution of the 159 sampling sites across Switzerland used in our study. *Alainites muticus* sex ratios significantly vary between populations (*GLM*, *p<0.001*). Populations are colored according to population sex ratio, given as the proportion of nymphs large enough to be sexed which are female (*e.g*., populations with only females are represented in red).

### Parthenogenetic capacity

Parthenogenetic reproduction is expected to generate female-biased sex ratios, with a more extreme female-bias in populations with a higher frequency of parthenogenesis. To determine whether the parthenogenetic capacity of females is positively correlated with population sex ratios, we aimed to obtain virgin females from 20 of the 159 *A. muticus* populations, by rearing female nymphs in the absence of males. Populations were chosen to cover the range of population sex ratios found in *A. muticus* (Fig. 1). Up to 20 late instar female nymphs were collected from 16 of the 20 populations (during May-July 2016, May-June 2017 and April-May 2018), no late instar nymphs could be obtained from the four remaining populations. Female nymphs were taken to the laboratory and reared to adulthood in aquariums in a climate chamber (water temperature: 12±1°C, room temperature: 22±2°C, relative humidity: 50%, 12L:12D photoperiod). Aquariums were supplied with lake water, in order to provide nutrients for nymphal growth until the final molt. The water temperature was maintained at 12°C (±1°C) using a thermoregulator and continuous water flow. Oxygenation was supplied through the water flow as well as from bubblers placed in each aquarium. Nymphs were reared in partially immersed floating cages (Supplementary figure S1). Submerged parts were surrounded by thin mesh allowing water flow while preventing mayflies from escaping, and the aerial parts allowed individuals to emerge. Emergences were checked daily, and subimagoes were transferred to individual cages. Once they completed their final molt (from subimago to imago), we allowed them to lay (unfertilized) eggs into a Petri dish (Ø 55 mm) filled to ¾ with filtered (0.2 μm) lake water. Using these conditions, we obtained 56 virgin *A. muticus* females, from which 11 did not lay any eggs spontaneously. For these females, we extracted the eggs by dissecting their abdomen after they died (within 0-12 hours). We counted the number of eggs per clutch, divided them into two to three new Petri dishes (Ø 35 mm) to reduce egg densities per dish and facilitate observation of hatchlings, and maintained them at 10°C with a 12L:12D photoperiod. After about three weeks of incubation, the eggs began to hatch. Hatchlings were counted and removed from each Petri dish every day, and the cumulative number of hatched eggs was recorded for each virgin female. After the cessation of hatching, we calculated the hatching success (proportion of unfertilized eggs that hatched) for each female. The population-level parthenogenetic capacity was estimated as the mean hatching success across all females collected from that population. In total, we obtained one to seven egg clutches (*i.e*., eggs from 1-7 females) per population.

### Statistical analyses

We first determined whether there was significant variation in sex ratios among sites using a binomial Generalized Linear Model (GLM) in R v.3.6.2 (R Development Core Team, 2019).

We then tested whether the parthenogenetic capacity in each population (measured as the mean hatching success of unfertilized eggs) affects the population sex ratio. To do so, we ran a quasibinomial GLM (generalized linear model with binomial error distribution, corrected for overdispersion) followed by an *F-test* from the ‘car’ package v.3.0-6 (Fox & Weisberg, 2019). Data points were weighted in the analysis according to sample size (the number of females tested per population).

Finally, in order to identify ecological correlates of female-biased sex ratios (*i.e*., frequency of parthenogenesis), we used Structural Equation Modeling (SEM) which tests for the nature and the magnitude of direct and indirect effects of each explanatory variable on the response variable (*e.g*., Grace *et al*., 2010; Eisenhauer *et al*., 2015; Fan *et al*., 2016). Specifically, we tested whether *A. muticus* population density, macro-invertebrate community diversity, and altitude significantly explained sex ratio variation across populations. Based on hypothesized causal relationships and correlations among the variables in the SEM, we built our initial meta-model (Fig. 2) and developed a structural equation model (Fig. 4, see results) that was analyzed using the ‘piecewiseSEM’ package v.2.1.0. A SEM is built using a list of structured equations, which can be specified using common linear modeling functions in R, and thus can accommodate non-normal distributions, non-independence of observations, etc. (Lefcheck, 2016). Then, based on model fit indices, calculated for the overall goodness of fit (*i.e*., Fisher’s C, cfi) and for each path (*i.e*., p-value and standard error), we evaluated model-data consistency to determine whether there were missing links in the initial meta-model, as well as to determine the support for tested links. The Fisher’s C statistic tests the hypothesis that there is a discrepancy between the model-implied covariance matrix and the original covariance matrix. Therefore, a non-significant discrepancy (p>0.05) indicates an acceptable model fit. The comparative fit index (cfi) represents the amount of variance that has been accounted for in the covariance matrix. A higher value indicates a better model fit (best is >0.95). We used these two indices as well as AIC from the ‘AICcmodavg’ package v.2.2-2 (Mazerolle, 2019) for model selection. In the final model, we corrected the model outputs for spatial autocorrelation using Moran’s I (Moran, 1950) within the ‘ape’ package v.5.3 (Paradis & Schliep, 2018).

**Figure 2.**
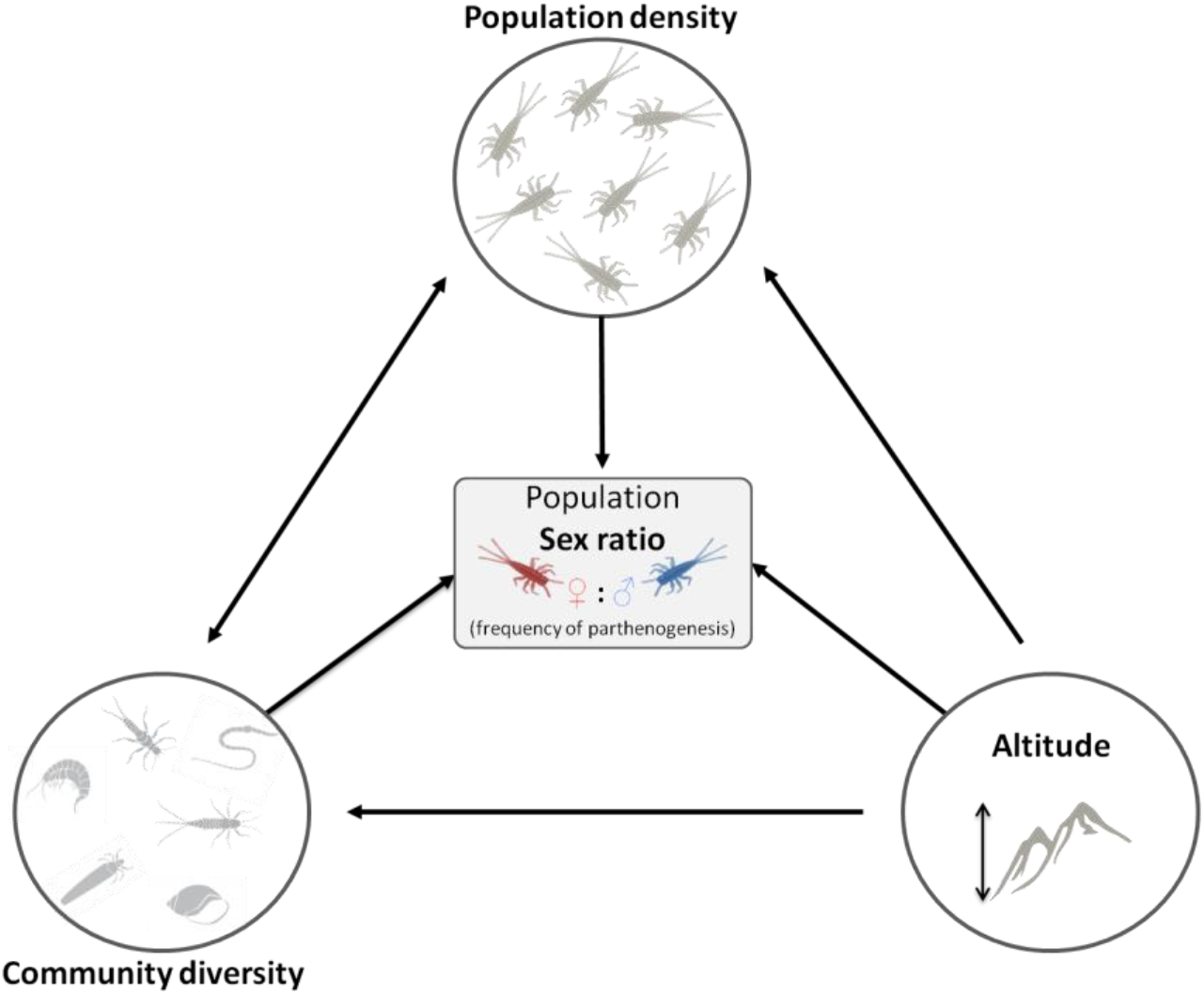
**Structural equation meta-model showing interconnections between ecological variables** (*i.e., Alainites muticus* population density, macro-invertebrate community diversity and altitude) **and population sex ratios** (used as a proxy for the frequency of parthenogenesis). Black arrows represent expected direct effects between two variables.

Finally, in order to test whether population variables were relatively stable or fluctuate over time, we developed a new structural equation model (Fig. 6, see results), using the data available for both surveys (*i.e*., 92 populations).

All plots were generated using the ‘ggplot2’ package v.3.3.5 (Wickham, 2016), in addition to the ‘rworldmap’ package v.1.3-6 (South, 2011) for the map in Fig. 1.

## Results

### Field evidence of parthenogenetic reproduction

In order to determine whether natural populations with a higher proportion of females also feature a higher parthenogenetic capacity, we used the average level of hatching success of unfertilized eggs of all 16 surveyed populations and checked whether it was related to the population sex ratio. In total, we were able to obtain unfertilized eggs from 56 females (one to seven females per population) and 51’771 eggs were observed for their parthenogenetic development (~1’000 eggs per female, range 372-2764). Hatching successes of unfertilized eggs (averaged across females) significantly affected population sex ratios (Fig. 3; *GLM, r* = *0.90*, *p*<*0.001*), meaning that a high parthenogenetic capacity of females in the field likely translates into female-biased population sex ratios. Given this result, we used sex ratios as a proxy of parthenogenetic capacity in natural populations.

**Figure 3.**
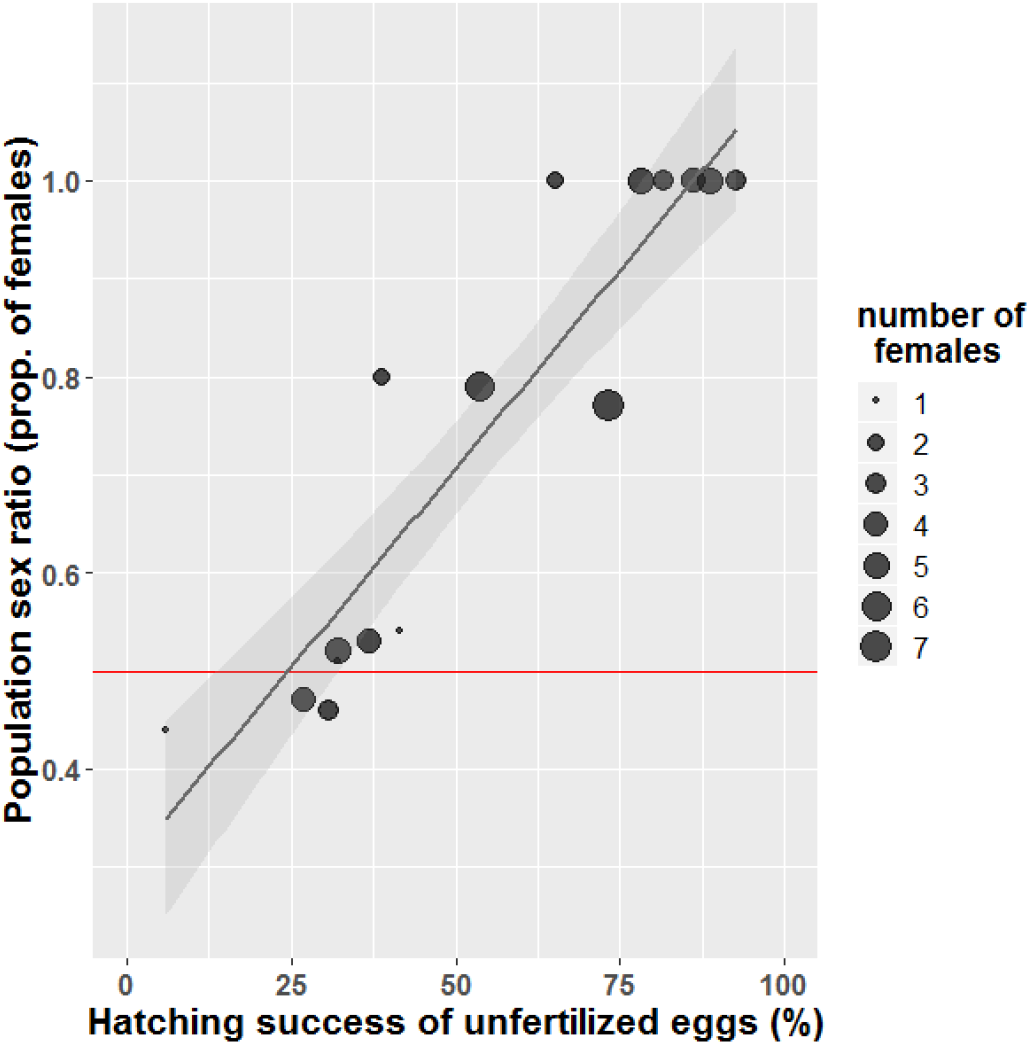
Parthenogenesis likely generates female-biased sex ratios in the field. Dot sizes are proportional to the number of females used to measure the parthenogenetic capacity for each population (*i.e*., one to seven females). Population sex ratios were estimated based on 15-322 sexed individuals per population (mean=90.4, median=54). The black line represents the fitted linear model (GLM, *r* = *0.90*, *p*<*0.001*), and the grey shadow represents the 95% confidence interval on the fitted values. The horizontal red line represents a balanced sex ratio with equal numbers of males and females.

### Densities and mate limitation

Population densities of the 159 *A. muticus* sites ranged from 15 to 450 sexed individuals per sampled surface (mean=77.4, median=52). Our SEM revealed that population densities have a significant direct and negative effect on population sex ratios (the proportion of females) (Figures 4 and 5A, *r* = −*0.20*, *p*=*0.002*), suggesting that mate limitation (or reduced harassment from males) selects for increased parthenogenetic capacity in natural *A. muticus* populations (see also Supplementary figure S2).

**Figure 4.**
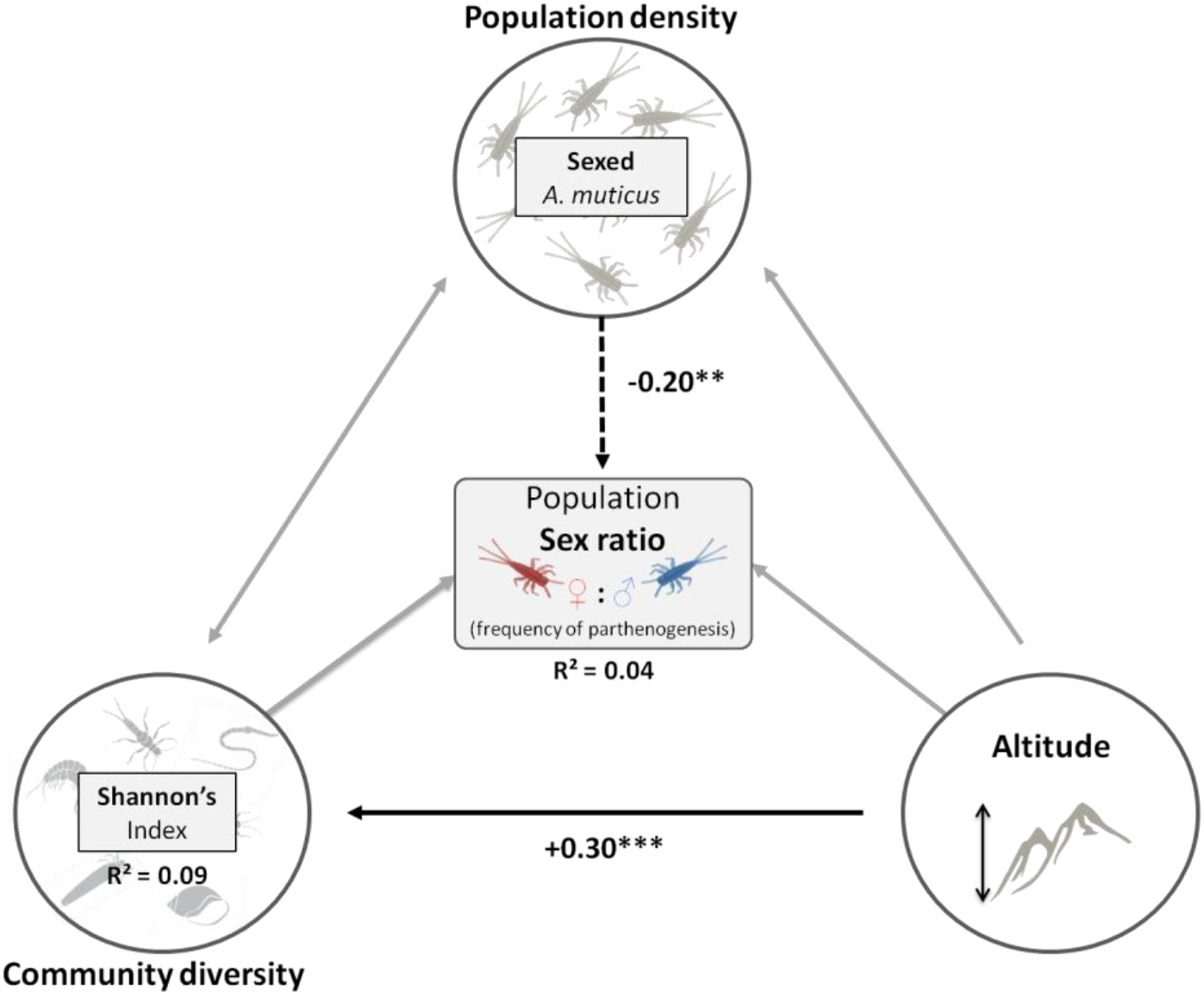
Structural equation model pathways fitted to collected data. Black arrows represent significant paths with positive (solid line) or negative (dashed line) effects (*p*<*0.05*). Grey arrows represent non-significant paths that were removed from the model (*p*>*0.30*). Path coefficients correspond to standardized effects, and R^2^ values are displayed on response variables, representing the proportion of variance explained. Test statistic *F* = 5.61, with 6 degrees of freedom, p-value = 0.47 and cfi = 1 (indicating close model-data fit, see Material & Methods for details).

**Figure 5.**
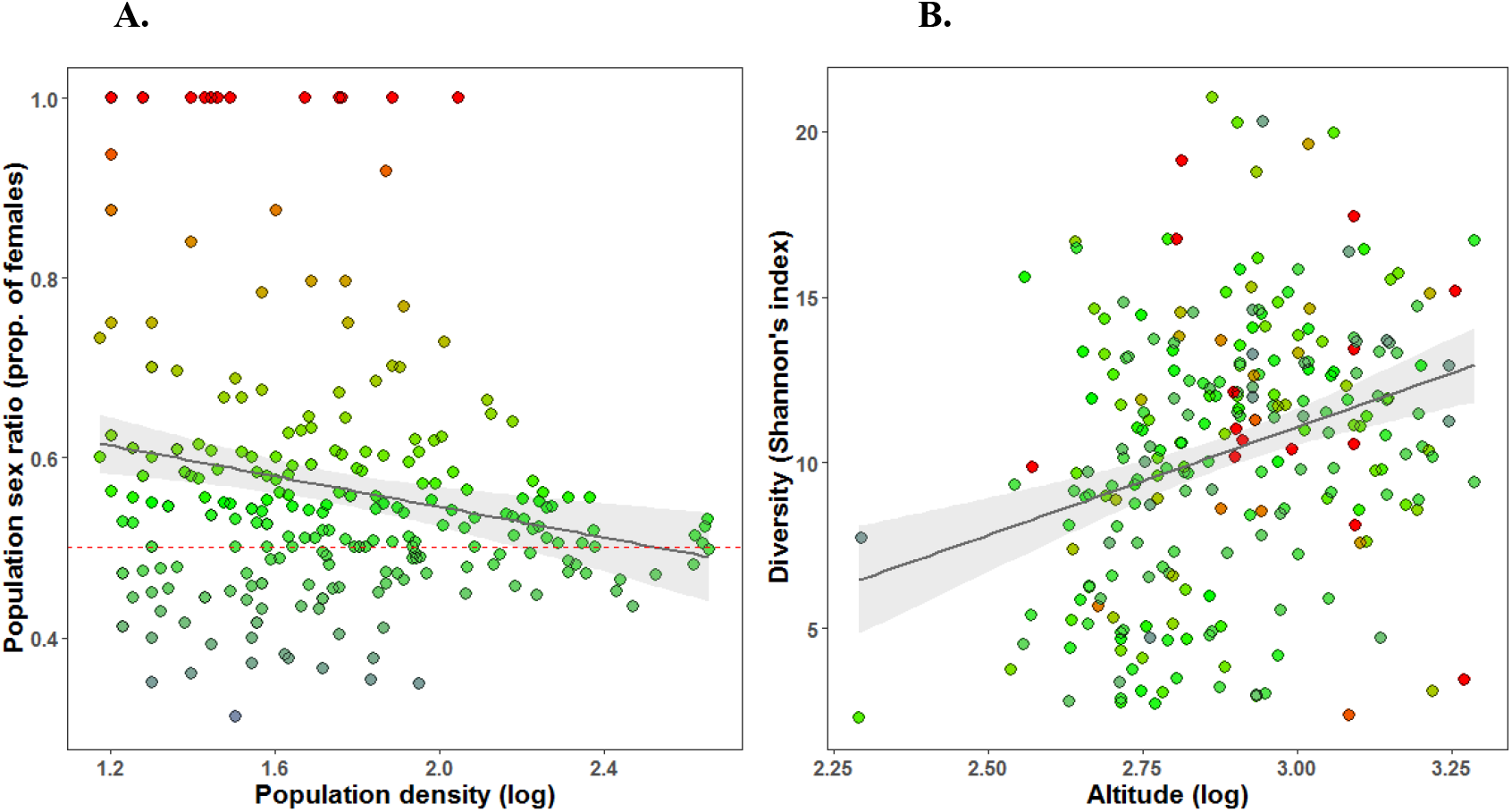
Multivariate partial plots for the significant paths in Figure 4. Note that for the 92 out of the 159 populations that were surveyed twice, both values are plotted (site ID as a random factor in the model). The partial plots for the non-significant paths are in Supplementary figure S4. The color gradient shows the population sex ratios (proportion of females), as in Fig. 1. The black line represents the fitted linear model and the grey shadow represents the 95% confidence interval on the fitted values. **A)** Partial relationship of the direct significant effect of population densities on population sex ratios. The horizontal red dashed line represents a balanced sex ratio (see also Supplementary figure S2). **B)** Relationship between altitude and community diversity (see also Supplementary figure S3). Note that the positive effect of altitude on community diversity was for the altitudinal range of *A. muticus* populations. Across the full altitudinal range of all surveyed sites (including those without *A. muticus*), diversity peaks at mid elevation (see Supplementary figure S5).

### Community diversity and competition hypotheses

According to the Red Queen, the Tangled Bank and Sib Competition hypotheses, we expected sexual reproduction to be more prevalent in more complex and diverse communities because of fast co-evolutionary dynamics and high levels of competition. Using Shannon’s index as an indicator of community diversity within the SEM (see Material & Methods for details), we did not find any direct effect of community diversity on *A. muticus* sex ratios (*i.e*., parthenogenesis frequencies), resulting in the removal of this non-significant path from the model (Fig. 4 and S4A). In addition, we did not find any indirect effect of community diversity on sex ratios (Fig. 4 and S4). Thus, our analyses do not detect an effect of community diversity on selection for sex in natural populations, neither directly nor indirectly. Note that similar results are obtained using other diversity indexes such as Margalef’s index, Simpson’s index or Hill’s numbers (results not shown).

### Altitude and geographical parthenogenesis patterns

Sites differ in various abiotic characteristics, and frequencies of parthenogenesis could be affected by those abiotic factors. We focused on altitude within the SEM, because of the pattern frequently observed in among species comparisons, where asexual lineages are more likely to be found at higher altitudes than their sexual counterparts (*i.e*., geographical parthenogenesis). However, we found no direct effect of altitude on population sex ratios (Fig. 4 and S4B), which does not corroborate the pattern observed in among species comparisons (Tilquin & Kokko, 2016.). We also did not find any indirect effect of altitude on sex ratios (Fig. 4 and S4). Nevertheless, there was a significant direct and positive effect of altitude on community diversity (Fig. 4 and 5B, *r* = *0.30*, *p*<*0.001*, see also Supplementary figure S3), which means that at higher altitudes, aquatic communities are more diverse. Note that similar results are obtained using other diversity indexes such as Margalef’s index, Simpson’s index or Hill’s numbers.

### Temporal variability of populations

In order to assess the fluctuation of population variables over time, we ran a new SEM with the 92 (out of the 159 populations) which were surveyed twice (Fig. 6). We found that sex ratios remained relatively stable between surveys, as shown by the significant positive correlation between sex ratios of the two surveys (Fig. 6 and 7A, *r* = *0.65*, *p*<*0.001*). This result suggests that sex ratios and thus the frequency of parthenogenesis within populations are relatively stable (41% of the variance explained). Most variation in sex ratios between years was for populations with low densities, which is most likely due to sampling effects, since sex ratio estimates are less exact for small sample sizes. Population densities also remained relatively stable between surveys, as shown by the significant positive correlation between densities of the two surveys (Fig. 6 and 7B, *r*=*0.21*, *p*=*0.04*). However, there was considerably more variation than for sex ratios, with only 5% of the variance explained (Fig. 6). Similarly, we also found that community diversities of the two surveys were significantly correlated (Fig. 6 and 7C, *r* = *0.23*, *p*=*0.04*), suggesting that diversities are also relatively stable over time, with 10% of the variance explained (this correlation is stronger if considering all the 500 sites, *i.e*., including sites without *A. muticus* present, *r* = *0.45*).

**Figure 6.**
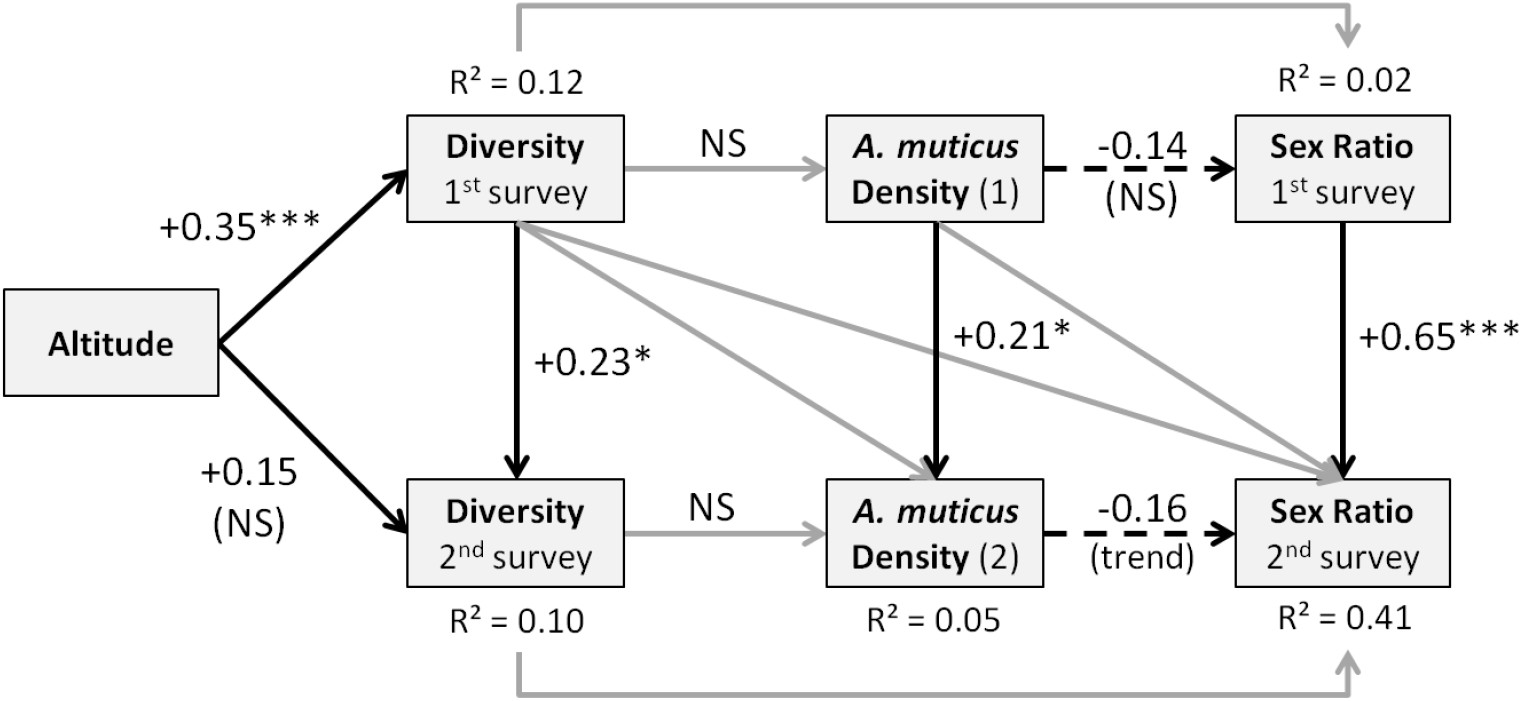
SEM including two surveys per site (n=92) to consider potential temporal fluctuation of population variables. Gray arrow: non-significant paths that were removed from the initial model. Dashed black arrows: negative effects. Solid black arrows: positive effects. R^2^ values are displayed on response variables, representing the proportion of variance explained. Test statistic F = 31.37, with 26 degrees of freedom, p-value = 0.22 (indicating close model-data fit, see Material & Methods for details).

**Figure 7.**
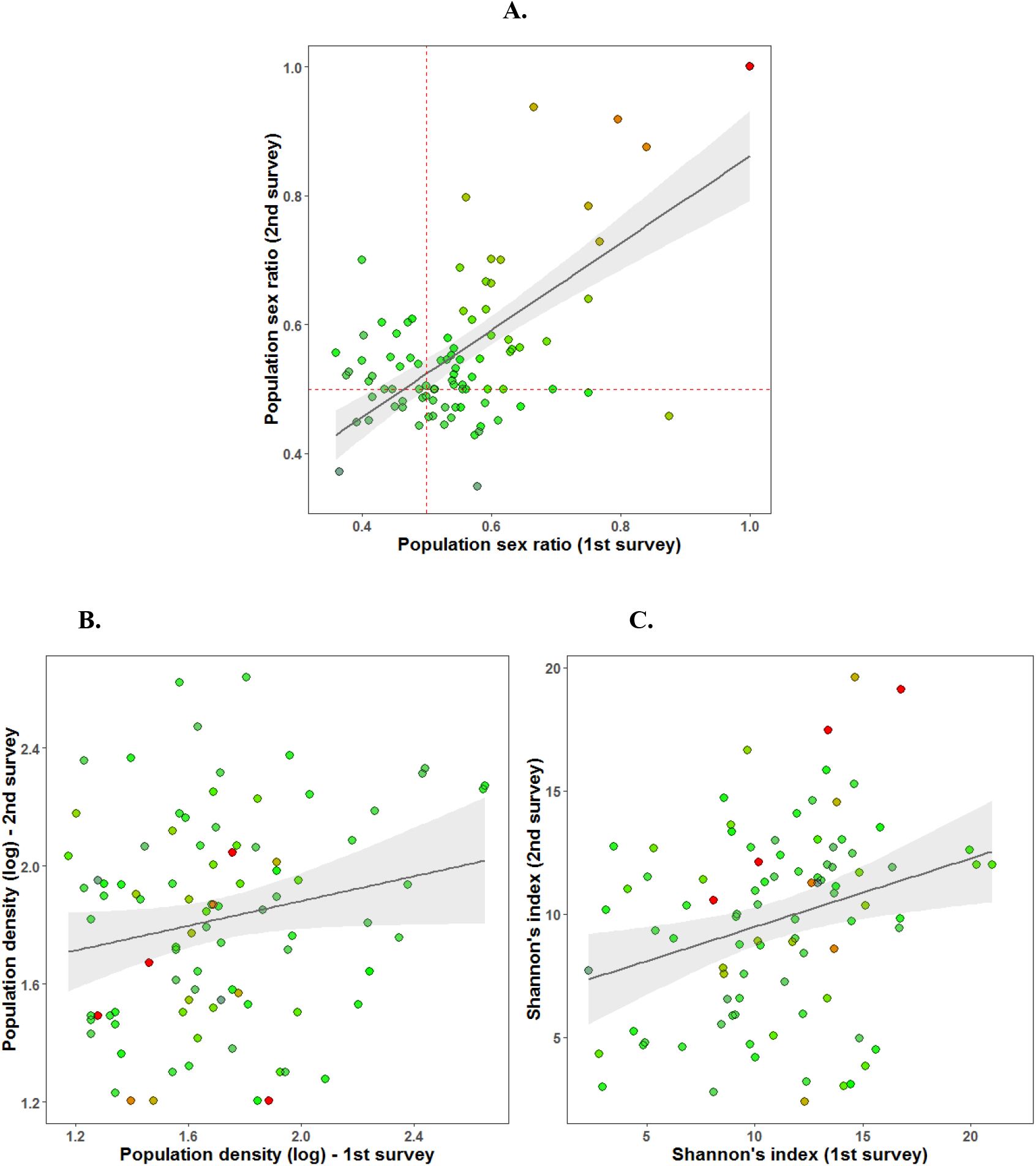
Multivariate partial plots over time. Significant positive effect of the first survey on the second one for population variables. The black line represents the fitted linear model and the grey shadow represents the 95% confidence interval on the fitted values. **A**) Population sex ratios (the red dashed lines represent the balanced sex ratios, *r* = *0.65*, *p*<*0.001*), **B**) *A. muticus* population densities (estimated from the number of sexed individuals, *r* = *0.21*, *p*=*0.04*), and **C**) Macro-invertebrate community diversities (*r* = *0.23*, *p*=*0.04*). The color gradient shows the population sex ratios (proportion of females), as in Fig. 1.

Note that we did not detect any crossed effect of variables from the first on the second survey. Notably, population density of the first survey does not affect the sex ratio of the second survey, meaning that sex ratios are apparently not shifted in the short term according to the past local densities, as would be expected if parthenogenetic capacities were plastic. In order to further investigate this, we looked at the within population variation between surveys for sex ratios and densities (Fig. 8).

**Figure 8.**
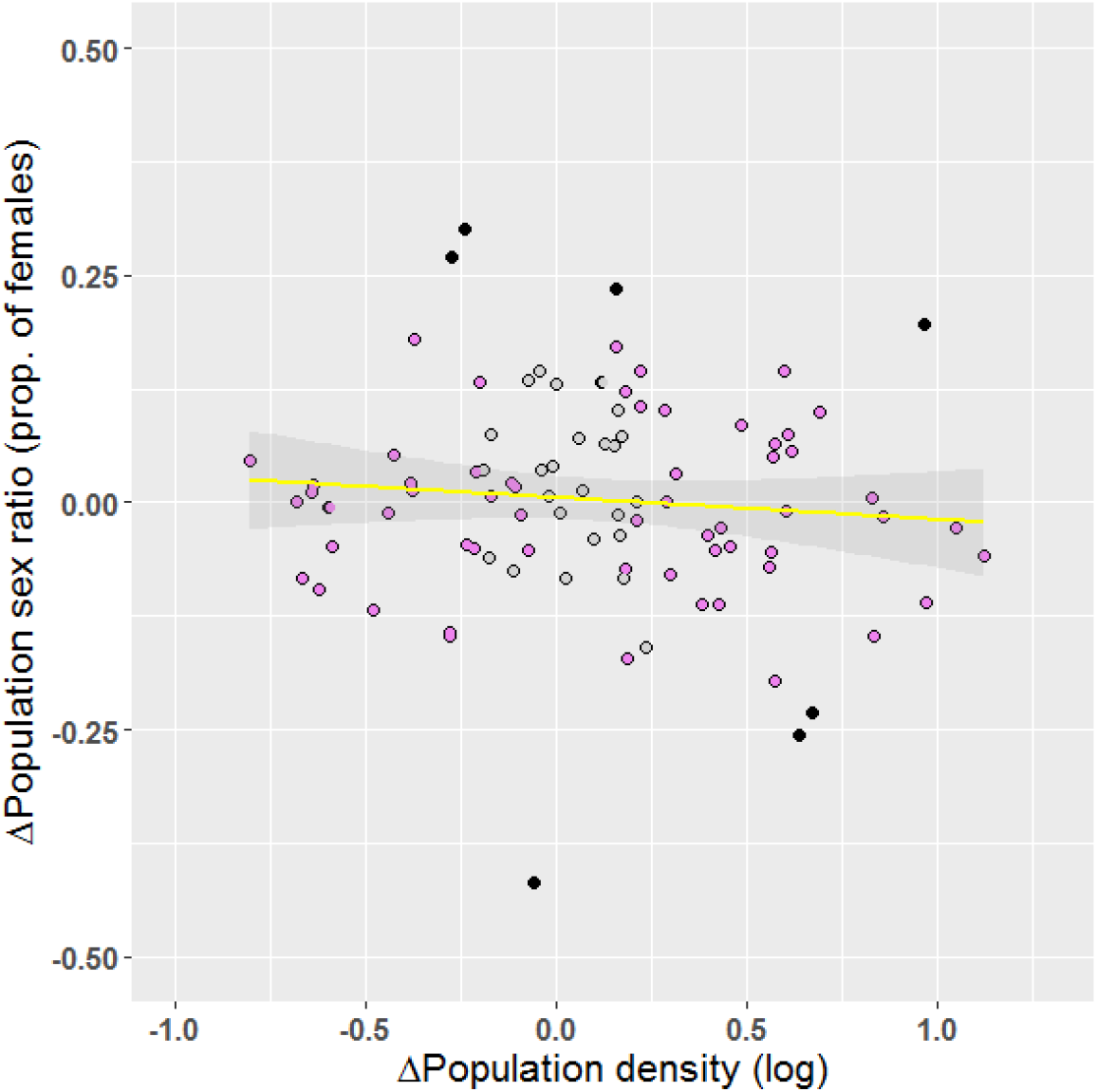
Within population differences between surveys for sex ratio and density. The yellow line shows the overall relationship (n=92). The black dots show populations that display a significant change in sex ratio over time (n=7). The pink dots show populations that display a significant change in density over time (n=65).

We found no significant correlation between the extent of the within population density shifts and the within population sex ratio shifts between surveys (Fig. 8, *p*=*0.38*), again suggesting that sex ratios are not shifted in the short term according to the current local densities. However, some populations displayed a significant change in sex ratio (n=7, Fig. 8), and most populations displayed a significant change in population densities (n=65, Fig. 8). For the latter, the change in density was not correlated with the change in population sex ratio (Fig. 8, pink dots), as expected if parthenogenetic capacities were not very plastic.

## Discussion

The aim of our study was to investigate how different ecological contexts influence reproductive strategies in the facultatively parthenogenetic mayfly *Alainites muticus*. We found a positive relationship between the parthenogenetic capacity of females (population average hatching successes of unfertilized eggs) and population sex ratios (proportion of females), revealing the use of parthenogenesis in natural populations of this species (90% of the variance explained), and therefore showing that population sex ratios can serve as a proxy of parthenogenetic capacities within natural populations. However, parthenogenesis is apparently only rarely used, given the overall high percentage of natural populations with balanced sex ratios in our survey (79.2% of the 159 sampling sites). This suggests that females generally prefer to reproduce *via* sex than parthenogenesis, or that if females have mated, they are unable to produce parthenogenetic eggs.

We found that the use of parthenogenesis is more widespread in populations where the density of individuals is low, as revealed by a significantly negative effect of *A. muticus* densities on population sex ratios, even though the explained variation among populations is small (only 4%, Fig. 4). Two different mechanisms could increase the frequencies of parthenogenesis in low-density populations. First, parthenogenesis could be used for reproductive assurance when females are mate limited. Mate limitation is expected to be widespread in mayflies as they are characterized by a very short adult lifespan, which may generate strong selection for reproductive assurance (Liegeois *et al*., 2021). Consistent with this idea, negative correlation between sex ratios and/or hatching successes of unfertilized eggs and population densities have also been reported for the mayfly species *Eurylophella funeralis* (Sweeney & Vannote, 1987), *Ephemerella notata* (Glazaczow, 2001), *Ephoron shigae* (Tojo *et al*., 2006) and *Stenonema femoratum* (Ball, 2002). Reproductive assurance through facultative parthenogenesis can generate more strongly female-biased sex ratios, which in turn increases mate limitation for females and selection for parthenogenesis in a positive feedback loop, which can result in the loss of males (Schwander *et al*., 2010).

Widespread facultative parthenogenesis and selection for reproductive assurance could thus also help explain why mayflies have more known parthenogenetic species than any other animal order (Liegeois *et al*., 2021). A second mechanism that can generate negative correlation between population densities and parthenogenetic frequencies is costly harassment from males (Kawatsu, 2013; Gerber & Kokko, 2016; Burke & Bonduriansky, 2017). Specifically, females with facultative parthenogenesis may pay the cost of resisting male harassment and mating attempts if they try to reproduce parthenogenetically. Thus, parthenogenesis would be more costly to females in populations with high frequencies of males. In this case, females reproducing sexually will be favored at high densities as harassment would likely increase with male availability (Gerber & Kokko, 2016). Notably, male coercion might also explain why parthenogenesis is only rarely used in our surveyed natural populations with facultatively parthenogenetic females.

Regarding community diversity, the Tangled Bank and Sib Competition hypotheses suggest that sexual reproduction should be favored under intense inter- and intra-specific competition in stable heterogeneous environments, where relatively saturated communities lead to a continuous and intense competition for resources (*e.g*., Bell, 1982; Scheu & Drossel, 2007). The Red Queen hypothesis proposes that sexual reproduction is favored because it can speed up co-evolution with parasites (*e.g*., Hamilton, 1980). Regarding these hypotheses, we investigated whether sexual reproduction of *A. muticus* is favored in more complex communities. However, we found no direct or indirect influence of community complexities (*i.e*., macro-invertebrate family diversities) on population sex ratios. A potential explanation for why we did not find the expected effect of community complexity is that our measures of community diversity did not take into account microorganisms (*i.e*., the most likely parasite communities) or big predators (*e.g*., fishes). In addition, parasites are a more important threat in high-density populations (*e.g*., Arneberg *et al*., 1998; Lagrue & Poulin, 2015), and population densities can also affect the level of intra-specific competition for resources. Because about 95% of a typical mayfly life cycle occurs at the nymphal stage and because adults do not feed, competition for resources is primarily a function of nymph densities. This might be another explanation for why we did not find the expected effect of community diversity on reproductive strategies, but why females in high density populations generally reproduce *via* sex instead of parthenogenesis in *A. muticus*.

As for altitude, we expected a positive relationship with the frequency of parthenogenesis which we did not find in *A. muticus*. Indeed, according to the patterns observed in geographical parthenogenesis, asexual lineages are more likely to be found at higher altitudes than their sexual counterparts (reviewed in Tilquin & Kokko, 2016). Thus, it seems that altitude may affect the relative success of sexual and asexual species, but it does not influence the frequency of reproductive modes in facultative parthenogens (*i.e*., within species). A probable explanation for this difference is that the altitudinal pattern is driven by species-specific traits that are confounded with reproductive modes. Indeed, in geographical parthenogenesis (*i.e*., in between species comparisons), asexuals are often polyploid or hybrids, meaning that the pattern observed might not be caused by the reproductive mode *per se* (Kearney, 2005). In our survey, we further detected a positive effect of altitude on community diversity for sites where *A. muticus* was present. This result suggests that abiotic factors such as altitude could create situations presenting suitable habitat conditions for increased community diversities and potential effects on population sizes we did not detect in our dataset.

Finally, to investigate whether differences in population sex ratios were more likely due to plasticity or genetic differences between populations, we quantified the fluctuation of population variables over time, using the 92 populations that were surveyed twice. We found that population sex ratios do not vary extensively over time. Population densities also remained relatively stable, even though they varied much more than sex ratios between surveys. We also found no significant correlation between the extent of the within population density shifts and the within population sex ratio shifts. In combination, these results suggest that sex ratios are not modified in the short term according to the current local densities, and are thus more likely generated by differences in parthenogenetic capacities between populations rather than plastic changes in the use of parthenogenesis.

Our population surveys also revealed an unexpected clustering of female-only populations of *A. muticus* south of the Alps, which could indicate the presence of obligately parthenogenetic lineages. Unisexual populations of *A. muticus* are also known in eastern Ukraine (Martynov, 2013), supporting the idea that this species might be characterized by reproductive polymorphism and geographical parthenogenesis. However, formally testing this idea, as well as investigating potential genetic differentiation between females with different reproductive modes, required different types of data and is a challenge for future studies.

To conclude, we suggest that low population densities tend to select for females with higher parthenogenetic capacities, likely because low densities generate situations of mate limitation for females or are associated with reduced costs generated by male coercion. The extremely short adult lifespan and limited dispersal abilities of mayfly may frequently generate locally low population densities and thus provide a steppingstone towards obligate parthenogenesis. Yet, even when females have the capacity to reproduce *via* parthenogenesis, they generally reproduce sexually, indicating that the benefits of sex for females must exceed its costs in most ecological situations.

## Acknowledgements

We are grateful to Nicolas Martinez and Markus Fluri (Hintermann & Weber AG, Reinach), who allowed us to use the material from the BDM surveys (which are financed by the Federal Office for the Environment) and who provided the ecological data for each sampling site. We would like to thank Jean-Luc Gattolliat for his help in the correspondence between Hintermann & Weber AG, the Museum of Zoology and the University of Lausanne. We also thank Victor Othenin-Girard for his help in separating *A. muticus* mayflies for the sex ratio calculations, as well as Darren J. Parker and Alexander Brandt for their comments on the manuscript. We thank James B. Grace for his helpful explanations on structural equation modeling. This work was supported by funding from the University of Lausanne and the Swiss National Science Foundation (grant numbers PP00P3_139013 and PP00P3_170627).

## Data and code availability

All data and code will be available in Dryad under https://doi.org/xx.xxxx/dryad.xxxxx.

