## Supplementary Materials for "What ecological factors favor parthenogenesis over sexual reproduction? A study on the facultatively parthenogenetic mayfly *Alainites muticus* in natural populations"

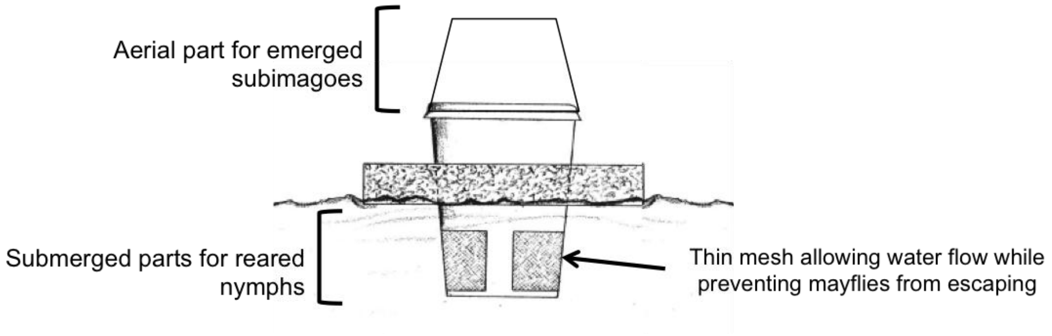


**Supplementary figure S1**. Partially immersed floating cage.

### Supplementary Materials

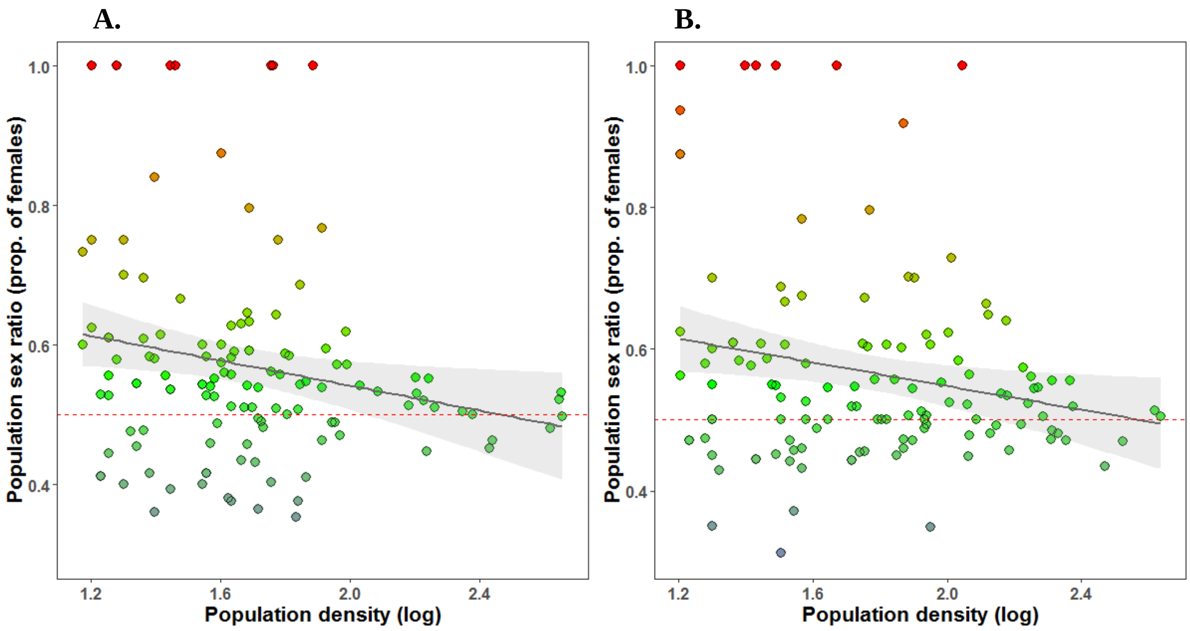


**Supplementary figure S2**. Significant negative effects of *A. muticus* densities on population sex ratios split between the first (**A.** n=126) and the second (**B.** n=125) survey. Population densities are estimated from the number of sexed individuals. The black line represents the fitted linear model and the grey shadow represents the 95% confidence interval on the fitted values. The color gradient shows the population sex ratios (proportion of females), as in Fig. 1. Note that 92 populations are common to both surveys.

### Supplementary Materials

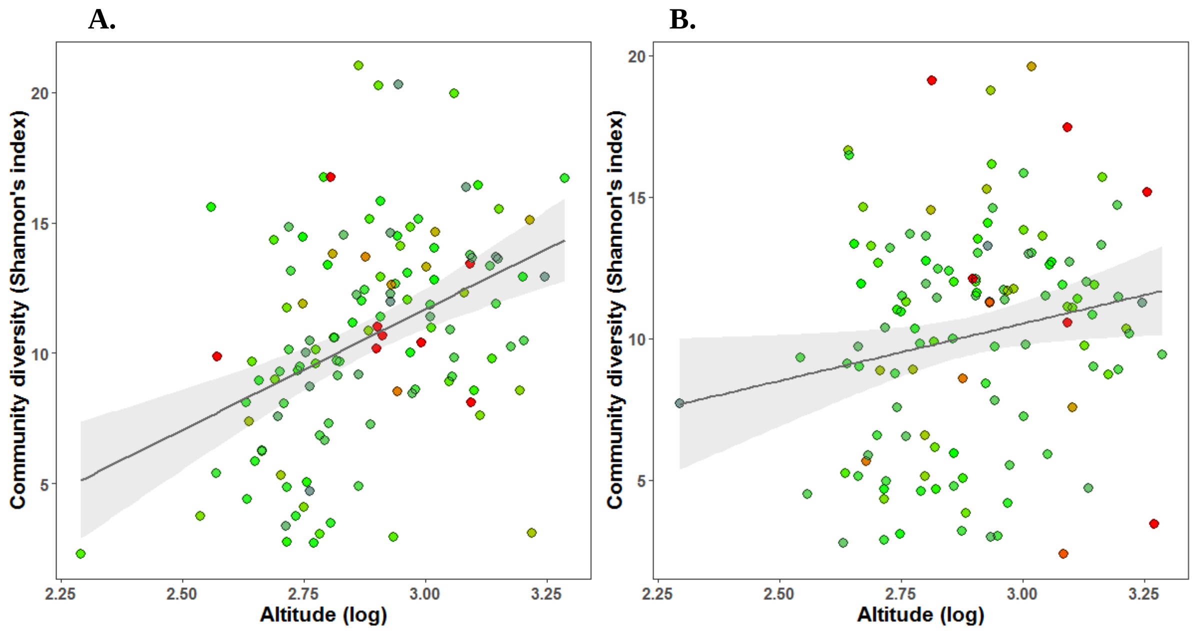


**<u>Supplementary Figure S3</u>**. Significant positive effects of altitudes on community diversities split between the first (Panel **A**, n=126) and the second (Panel **B**, n=125) survey. The black line represents the fitted linear model and the grey shadow represents the 95% confidence interval on the fitted values. The color gradient shows the population sex ratios (proportion of females), as in Fig. 1. Note that 92 populations are common to both surveys.

### Supplementary Materials

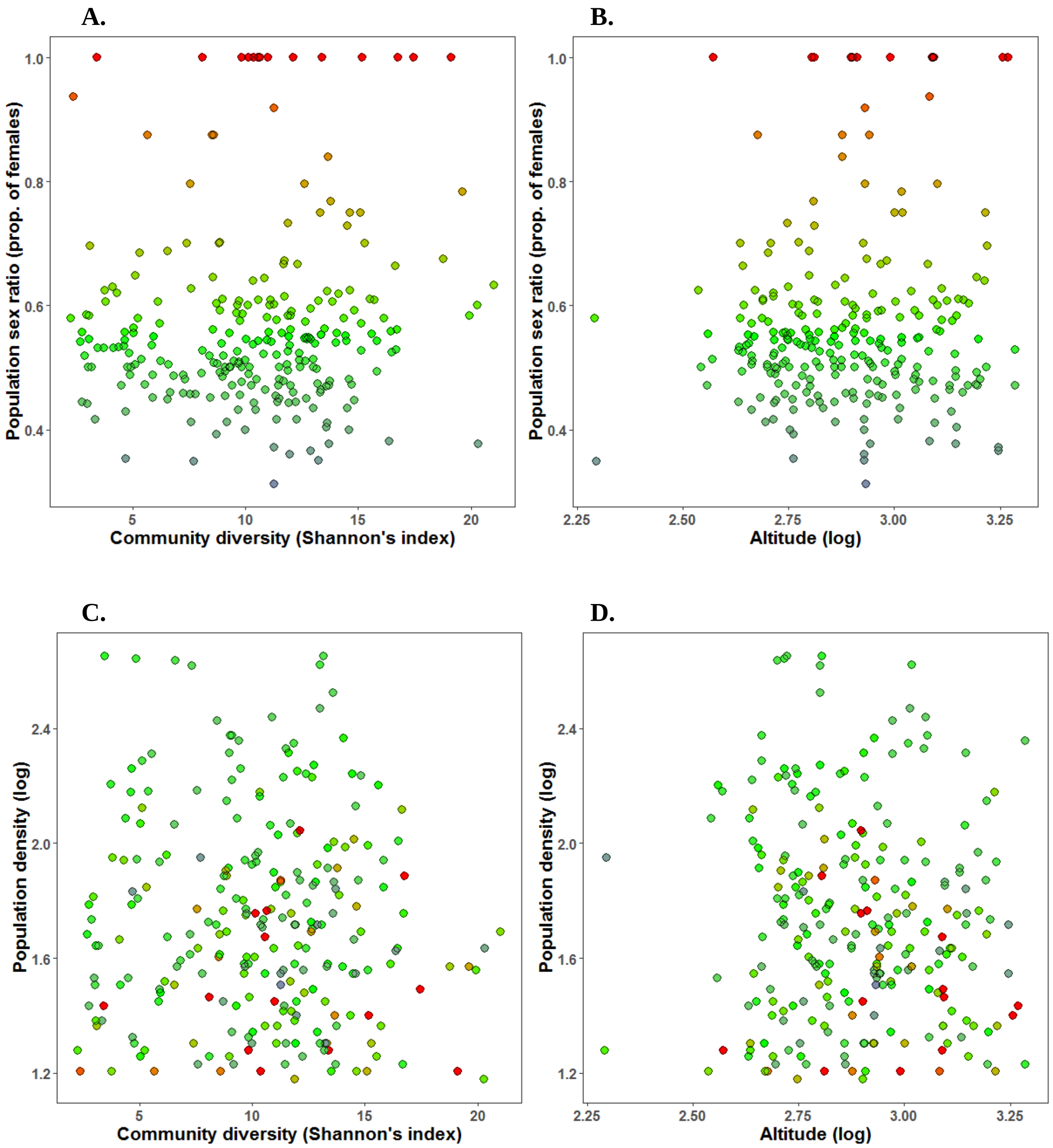


**Supplementary figure S4**. Plots for the non-significant paths in Figure 4. Relationships between community diversity and population sex ratio (**A**), altitude and population sex ratio (**B**), community diversity and population density (**C**), and altitude and population density (**D**). The color gradient shows the population sex ratios (proportion of females), as in Fig. 1. Note that the 92 out of the 159 populations that were surveyed twice, both values are plotted (site's ID as a random factor in the model).

### Supplementary Materials

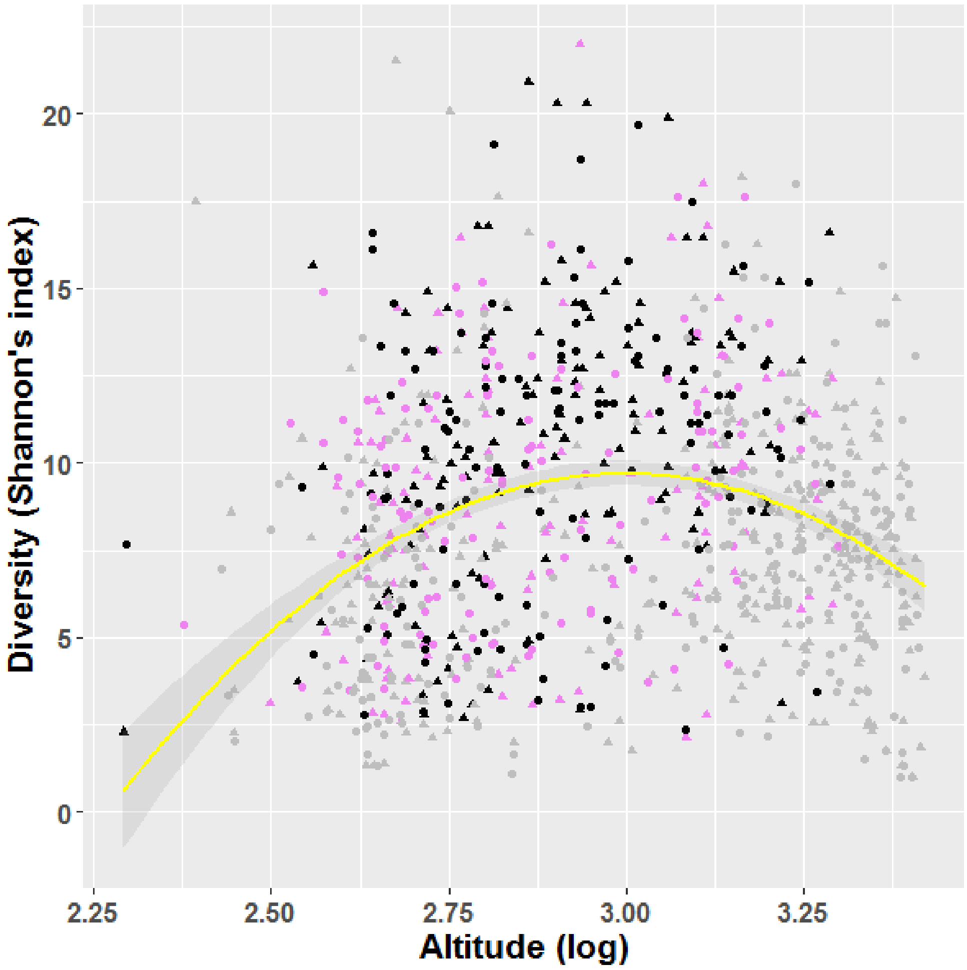


**<u>Supplementary figure S5</u>**. Diversity peaks at mid elevation. Black color = populations used within the SEM (*i.e.*, populations with >15 sexed *A. muticus*); purple color = populations with <15 sexed *A. muticus*; gray color = populations without *A. muticus*. Triangles = 1st survey (n= 490), circles = 2nd survey (n = 474).
